# Genome sequencing of the bacteriophage CL31 and interaction with the host strain *Corynebacterium glutamicum* ATCC 13032

**DOI:** 10.1101/2021.02.12.430922

**Authors:** Max Hünnefeld, Ulrike Viets, Vikas Sharma, Astrid Wirtz, Aël Hardy, Julia Frunzke

## Abstract

In this study, we provide a comprehensive analysis of the genomic features of the phage CL31 and the infection dynamics with the biotechnologically relevant host strain *Corynebacterium glutamicum* ATCC 13032. Genome sequencing and annotation of CL31 revealed a 45-kbp genome composed of 72 open reading frames, mimicking the GC content of its host strain (54.4 %). An ANI-based distance matrix showed the highest similarity of CL31 to the temperate corynephage Φ16. While the *C. glutamicum* ATCC 13032 wild type strain showed only mild propagation of CL31, a strain lacking the *cglIR-cglIIR-cglIM* restriction-modification system was efficiently infected by this phage. Interestingly, the prophage-free strain *C. glutamicum* MB001 featured an even accelerated amplification of CL31 compared to the Δ*resmod* strain suggesting a role of cryptic prophage elements in phage defense. Proteome analysis of purified phage particles and transcriptome analysis provide important insights into structural components of the phage and the response of *C. glutamicum* to CL31 infection. Isolation and sequencing of CL31-resistant strains revealed SNPs in genes involved in mycolic acid biosynthesis suggesting a role of this cell envelope component in phage adsorption. Altogether, these results provide an important basis for further investigation of phage-host interactions in this important biotechnological model organism.

## 1. Introduction

The Gram-positive actinobacterium *Corynebacterium glutamicum* represents an important biotechnological platform strain used for the production of amino acids at million tons scale as well as for the production of organic acids, polymer precursors, and proteins [1]. This study focuses on derivatives of the model strain *C. glutamicum* ATCC 13032 (NCBI accession number BX927147) [2]. The genome of this strain includes three cryptic prophages, of which two (CGP1 & 2) are small and highly degenerated, while the third one (CGP3) was shown to be still inducible [3]. The CGP3 island also harbors a restriction modification (resmod) system (*cglIM*, *cglIR*, and *cglIIR*), which was previously shown to confer resistance to phage infection [4,5]. Besides the wild type strain *C. glutamicum* ATCC 13032, this study also includes two derivative strains: a strain lacking the resmod system (Δ*resmod*) and the prophage-free variant MB001 [6].

Bacteriophages represent a ubiquitous threat to biotechnological processes, since phage infection may lead to high biomass losses. Besides that, phages can serve as a source of important molecular biology tools and fully adapted modulators, which can be harnessed for metabolic engineering and synthetic biology applications [7,8].

Although the first phages infecting *C. glutamicum* were already described in the 1970s, until now the number of characterized phages infecting this host is limited and genome sequence is available for only few of them. Among those are the temperate phage Φ16 [9–11], as well as the virulent phages BFK20 [12,13], Φ673 and Φ674 [14], and P1201 [15]. In particular, the lytic phage BFK20 was well characterized with regard to its host spectrum and interaction, its transcriptional profile, and functions of single BFK20-encoded proteins [12,16,17]. Furthermore, BFK20 promoters were applied in an overexpression system for recombinant RNAs in *C. glutamicum* [18]. Another example of application of corynephage-derived systems is represented by Φ16. This temperate phage provided the basis for the construction of a site-specific integration vector for *C. glutamicum* [19].

This study focuses on the phage CL31, which was first identified in 1985 by Yeh and colleagues as a phage infecting C. *glutamicum* ATCC 15059 (previously known as *Corynebacterium lilium*) [20]. In further studies, Trautwetter and colleagues described CL31 as a siphophage with a narrow host range (4 out of 30 tested strains were lysed) consisting of approximately eight structural proteins and a linear 48 kb double-stranded DNA genome with cohesive ends [21]. Phage ‘Corlili’ listed in the actinobacteriophage database (phagesdb.org) actually refers to the very same phage [22].

In this study, we performed genome sequencing and annotation of phage CL31. Analysis of infection dynamics and transcriptome analysis confirmed the central role of the resmod system in phage defense and showed a significant impact of the presence of cryptic prophage elements on phage amplification. Single nucleotide polymorphisms (SNPs) conferring resistance to CL31 infection converged in pathways involved in mycolic acid synthesis, shedding light on the molecular adsorption mechanism of this CL31. Altogether, these data provide an important ground for further investigation of phage-host interactions in this important biotechnological model organism.

## 2. Materials and Methods

### 2.1 Bacterial strains, plasmids and growth conditions

Bacterial strains and plasmids used in this work are listed in Table 1. For cloning and plasmid construction the strain *Escherichia coli* DH5α was used. Unless stated otherwise, this strain was cultivated in lysogeny broth (LB, [23]) containing 50 μg/ml kanamycin. *Corynebacterium glutamicum* ATCC 13032 (Accession: BX927147) was used as a wild type strain [2]. This wild type strain and all derived *C. glutamicum* strains were cultivated in brain heart infusion medium (BHI, Difco Laboratories, Detroit, MI, USA), if necessary 25 μg/ml kanamycin was added. For the cultivation of *C. glutamicum*, a first pre-culture was inoculated with single colonies from agar plates either directly after transformation or after a streak-out of glycerol cultures. These pre-cultures were conducted in 4.5 ml BHI medium in test tubes or – depending on the purpose - in 1 ml BHI medium in 96-well deep well plates (DWPs) at 30°C overnight. For all infection experiments, a starting OD_600_ of 0.5 was used. If not stated otherwise, the multiplicity of infection (MOI) was set to 0.1. To start infection, bacteria and phages were mixed in 1/5^th^ of the final media volume and pre-incubated for 5 min at RT to allow phage attachment to the host cells. Subsequently, the culture was filled-up to the final volume and incubation was started at 30°C and 120 rpm. Additionally, growth experiments were conducted in the BioLector microbioreactor system (m2p labs, Baesweiler, Germany) [24]. Therefore, 750 μl BHI medium were inoculated to an OD_600_ of 0.5 in 48-well microtiter plates (Flowerplates, m2p labs). CL31 phages were added to reach the desired MOI and after a 10 min pre-incubation without shaking at RT, the main cultivation was conducted at 30°C and 1200 rpm shaking frequency. During this cultivation, backscatter was measured with an excitation wavelength of 620 nm (filter module: λEx/ λEm: 620 nm/ 620 nm, gain: 20) in 15 minutes intervals.

**Table 1.**
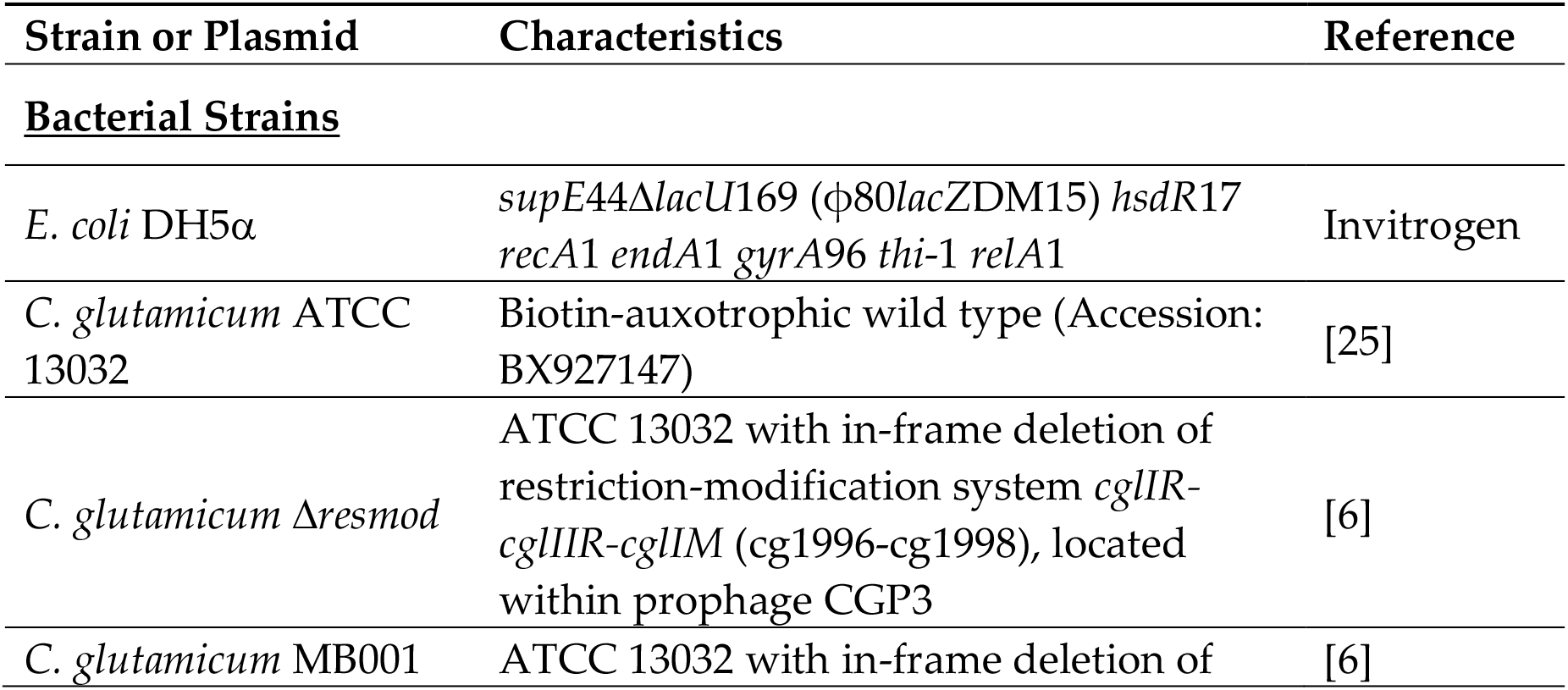
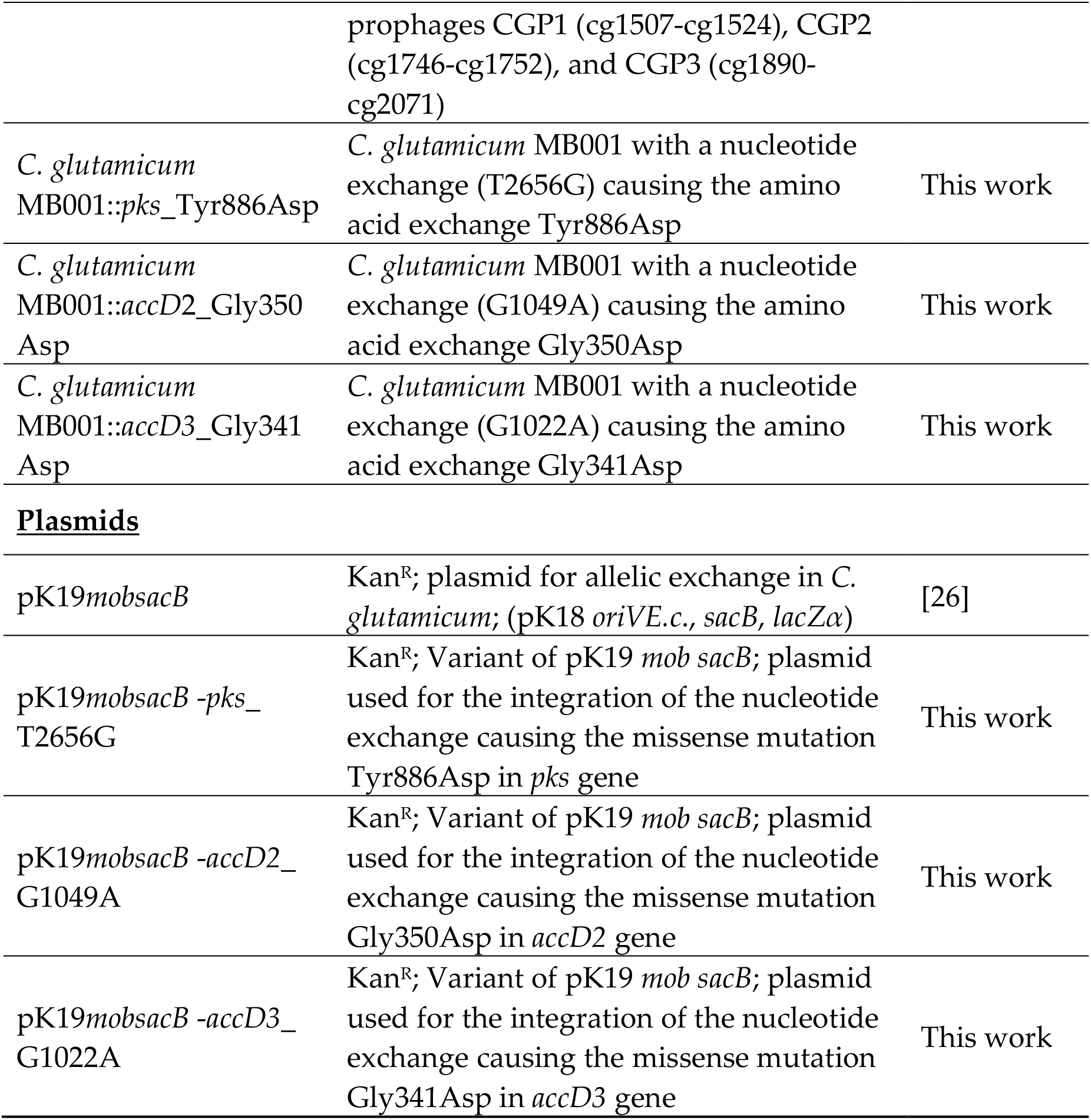
Bacterial strains and plasmids used in this study.

For double-layered agar overlay plates, a top layer of soft agar (0.4% agarose in BHI medium) containing the respective host bacteria (OD_600_ 0.5) was poured on top of a bottom agar layer (2% agar-agar in BHI medium).

### 2.2 Recombinant DNA work and construction of chromosomal insertions or deletions

All standard laboratory methods (PCR, DNA restriction, Gibson Assembly) were performed according to standard protocols and manufacturer’s instructions [23,27]. The used oligonucleotides as well as details regarding the plasmid construction are provided in the supporting information (Table S1 and Table S2). For genomic substitution of the mutated genes variants, a two-step homologous recombination system based on the suicide vector pk19*mobsacB* was used [28,29]. This vector contained 500 bps of each site flanking the targeted sequence in the genome of *C. glutamicum*. Successful re-integration was confirmed by colony-PCR and sequencing.

### 2.3 Phage Propagation

The initial phage stock of CL31 was provided by the Félix d'Hérelle Reference Center for bacterial viruses of the Université Laval (Québec, Canada). To amplify the phage, *C. glutamicum* MB001 was inoculated to an OD_600_ of 5 in 20 ml BHI medium, 7.5 μl CL31 lysate was added and the culture was incubated for 5 min at RT without shaking. Subsequently, the culture was filled-up to the final volume of 100 ml to reach the final cultivation OD_600_ of 0.5. This culture was then incubated at 30°C and 120 rpm shaking frequency. When the culture was lysed completely (~ 4 h), supernatants were harvested by centrifugation for 15 min at 4500 x g. The supernatants were sterile filtered (0.2 μm pore size), and the buffer was exchanged to sodium chloride/magnesium sulfate (SM) buffer (10 mM Tris-HCl pH 7.3, 100 mM NaCl, 10 mM MgSO_4_, 2 mM CaCl_2_) using Amicon Ultra Filter falcons with 30 kDa cutoff (MerckMillipore, Billerica, USA). The latter step was additionally used to concentrate the samples. Finally, the lysate was mixed with glycerol to a final concentration of 10% (w/v) and stored at −80°C until further usage.

### 2.4 Phage Purification with cesium chloride gradients

A further purification of the isolated phage particles was conducted as preparation for the proteomics measurements. For this purpose, cesium chloride (CsCl) was dissolved in SM buffer creating solutions of different densities (1.3, 1.4, 1.5, 1.6 g/ml). These solutions were subsequently layered into a centrifuge tube with 2 ml per density (with lowest density on top). 2 ml of pre-purified phage solution (in SM buffer) were added on top of this gradient and centrifuged in an ultracentrifuge at 150,000 x g and 4°C for 2 h.

During this centrifugation, a milky layer of phage particles appeared between the 1.3 g/ml and 1.4 g/ml layer. This milky layer was extracted, washed three times with SM buffer using Amicon Ultra centrifugal filters with 30 kDa cutoff (MerckMillipore, Billerica, USA) and subsequently tested for their phage titer.

### 2.5 DNA Isolation

The isolation of phage DNA was conducted as described by Hardy and colleagues [30]. To be more precise, 1 μL of 20 mg/mL RNAse A and 1 U/μL DNAse (Invitrogen, Carlsbad, CA, USA) were added to 1 mL of the filtered lysates to limit contamination by host nucleic acids. The suspension was incubated at 37° C for 30 min. Subsequently, EDTA, proteinase K and SDS were added to the mixture at final concentrations of 50 mM (EDTA and proteinase K) and 1% SDS (w/v), respectively. The digestion mixture was incubated for 1 h at 56 °C, before adding 250 μL of phenol:chloroform:isopropanol. The content was thoroughly mixed before centrifugation at 16,000 × *g* for 4 min. The upper phase containing the DNA was carefully transferred to a clean microcentrifuge tube and 2 volumes of 100% ethanol were added as well as sodium acetate to a final concentration of 0.3 M. After centrifugation at 16,000 x *g* for 10 min, the supernatant was discarded, and the pellet washed with 1 mL 70% ethanol. Finally, the dried pellet was resuspended in 30 μL DNAse-free water and stored at 4°C.

### 2.6 DNA Sequencing and Genome Assembly

The DNA library was prepared using the NEBNext Ultra II DNA Library Prep Kit for Illumina (New England Biolabs, Ipswich, MA, USA) according to the manufacturer’s instructions and sequenced using the Illumina MiSeq platform with a read length of 2 × 150 bp (Illumina, San Diego, USA). In total, 100,000 reads were subsampled, and de novo assembly as well as contig verification was performed using CLC genomics workbench 9.0 (QIAGEN, Hilden, Germany). The resulting genome displayed a ~4,600x average coverage.

### 2.7 Gene Prediction and Functional Annotation

Gene prediction and annotation was performed with slight modifications as described before by Hardy and colleagues [30]. Open reading frames (ORFs) in the phage genomes were identified using a combination of PHANOTATE v1.5.0 [31], GeneMark.hmm_PROKARYOTIC v3.26 [32] and the ORF prediction function of SnapGene software (from Insightful Science; available at snapgene.com). Functionally annotation was done using an automatic pipeline using multiphate v1.0 [33]. The functional annotation was automatically improved and curated using hidden Markov models (HMMs), and BLASTp [34] based similarity searches against different databases Prokaryotic Virus Orthologous Groups (pVOGs [35], NCBI viral proteins, and Conserved Domain Database CDD [36]), with the e-value cutoff 10−10. Additionally, BLASTp searches against NR database (e-value cutoff 0.05) and PhagesDB database [22] (e-value cutoff 1) were used to add broader information to the annotation table. The ends of the phage genome were determined with PhageTerm [37] using default parameters. The rearrangement of the CL31 genome based on the PhageTerm results showed an area that was not successfully resolved by the genome assembly. Thus, Sanger sequencing was used to further resolve this repeat-containing area (Figure S1). The annotated genome was deposited in GenBank under the accession number MW582634.

### 2.8 Genomic distance analysis

Genome-wide nucleotide k-mer frequency distribution was calculated across the complete > 2950 bacteriophage genomes using an approach based on machine learning in R [38,39]. Further details about code and methodology on https://bioinformaticshome.com/bioinformatics_tutorials/R/phylogeny_estimation.html.

The resulting frequency matrix was used to calculate the pairwise distance between the phage genomes with the Jensen-Shannon divergence method. The resulting distance matrix was used to display the genomic distance using a boxplot with ggplot2 package [40] in R.

### 2.9 Phage Classification

To establish the relationship between the newly sequenced CL31 phage with already known actinobacteriophage genomes, initially, we randomly selected the one representative phage genomes from each known group or cluster from the PhageDB [22]. A selected set of 31 phage genomes, including CL31, was used to calculate pairwise average nucleotide identity (ANI) using the python program pyani 0.2.9 [41] with the help of the ANIb method. The resulted average percentage identity matrix obtained from pyani was clustered and visualized using the R package ComplexHeatmap [42]. Additionally, pairwise intergenomic distances and similarities between the same set of Actinobacteriophage genomes were calculated using the VIRIDIC webserver [43].

### 2.10 Lifestyle prediction

We used a novel computational approach for actinobacteriophage lifestyle prediction based on counts of the set of specific genome-wide distributed conserved protein domain using a custom-designed R script. Initially, we identified the conserved protein domains from 2951 complete actinobacteriophage genomes against the CDD database [44] using RPSTBLASTN with e-value (0.001). The predicted protein domains from all these genomes were divided into two parts based on the known phage lifestyle (temperate and virulent) information. The list of protein domains specific to each corresponding lifestyle was collected and used as a base to classify new phage genomes. The first step in the classification process is identifying the protein domains from the individual phage genome (Ex: CL31) using RPS-TBLASTN. The identified protein domains were then mapped and counted against the base list of temperate and virulent specific domains. Finally, check the total number of the mapped particular protein domains within the individual genome and assign putative lifestyle based on the one which contains higher counts.

### 2.11 Phage Infection Curves and Titer Determination

The dynamics of CL31 infection were analyzed using bacterial growth (measured as OD_600_) and phage titer (measured with spot assays) for a time course of 24 h. Cultivations were conducted as described in 2.1. For shaking flask cultivations, an MOI of 0.1 was used and 1 ml samples were taken at the beginning of the cultivation (0 h), hourly up to 6 h and a final sample at 24 h after cultivation start. OD_600_ of these samples was measured and after a 5 min centrifugation at 16,000 x g and 4°C, the top 200 μl of the supernatant were diluted in SM buffer in a ten-fold series up to 10-8. Subsequently, 3 μl of each dilution were spotted on overlay agar plates with soft agar containing MB001 cells.

After an overnight incubation at 30°C, the phage titer could be determined by counting single plaques and calculating the plaque forming units (PFU) per ml.

### 2.12 Transcriptome Analysis (RNA-seq)

#### 2.12.1 Sample preparation and sequencing

For RNA sequencing, MB001 as well as WT cells were cultivated in shake flasks as described in 2.1. In addition to CL31-infected samples, we cultivated the strains also without CL31 as controls. Samples were extracted at 0, 0.5, 0.75, 1, 1.5 and 2 h after cultivation start (5 ml for RNA preparation, and 1 ml for OD_600_ measurement and phage titer determination). Phage titers were determined as described before and a time point before titer increase was chosen for RNA preparation. This preparation was conducted with the Monarch Total RNA Miniprep Kit (New England Biolabs) following the manufacturer’s instructions.

Subsequently, ribosomal RNA was depleted using NEBNext rRNA Depletion Kit (Bacteria) (New England Biolabs) with additional organism-specific spiked-in custom probes following the manufacturer’s protocols. RNA quality was determined before and after rRNA depletion using a TapeStation 4200 (Agilent Technologies Inc, Santa Clara, USA).

In a next step, the fragmentation of RNA, cDNA strand synthesis and indexing were carried out using the NEBNext® Ultra™ II Directional RNA Library Prep Kit for Illumina® (New England Biolabs) according to the supplier’s manual. The resulting cDNA was purified using Agencourt AMPure XP magnetic beads (Beckman Coulter, IN, USA) and libraries were quantified using the KAPA library quant kit (Peqlab, Bonn, Germany) and normalized for pooling. Pooled libraries were sequenced on a MiSeq (Illumina, CA, USA) using the MiSeq Reagent Kit v3 to generate single-end reads with a length of 150 bases. Base calling and preprocessing were conducted with the Illumina instrument software and stored as fastq output files.

#### 2.12.2 Bioinformatic transcriptome analysis

The quality of the sequencing reads was analyzed using FastQC (https://www.bioinformatics.babraham.ac.uk/projects/fastqc/). Because of good quality and absence of significantly over-represented (adapter) sequences, no further pre-processing was undertaken.

Further read analysis was conducted using CLC genomic workbench V. 10.1.1 software (QIAGEN, Hilden, Germany). The reads were mapped against the *C. glutamicum* genomes (Accession BX927147.1) and the CL31 genome and transcripts per million (TPM) values were calculated using the RNA-seq analysis tool of CLC genomics workbench using the following read alignment parameters: Mismatch cost: 2, Insertion cost: 3, Deletion cost: 3, Length fraction: 0.8, Similarity fraction: 0.8, Strand specific: both, Maximum number of hits for a read: 10. Subsequently, a TPM value table was exported and further processed by calculation of inter-replicate means and variances (expressed as a percentage of the relative difference between single replicate values). Furthermore, a fold-change was calculated based on each mean value.

### 2.13 Staining of CL31 for Fluorescence Microscopy

In order to visualize CL31 particles using fluorescence microscopy, 500 μl phage solution were mixed with 50 μl SYBR Gold (Invitrogen, Carlsbad, USA), diluted 1:1000 with SM buffer. After an incubation at 4°C in the dark for 1 h, the samples were transferred to an Amicon Ultra centrifugal filter falcon with 30 kDa cutoff (MerckMillipore, Billerica, USA). The sample were washed three times with 10 ml SM buffer, using centrifugation steps of 10 min and 4,000 rpm in a swinging rotor. After the washing steps, remaining 500 μl were collected and kept in a dark tube at 4°C until further use.

Bacterial cells were stained using SM buffer containing 100 ng/ml Hoechst 33342 and 250 ng/ml Nile red. For this purpose, a sample was taken out of an exponentially growing culture (OD_600_ ~3, cultivation as described before), washed with SM buffer and resuspended in the SM buffer containing Hoechst 33342 and Nile red. After a 15 min incubation at room temperature, cells were pelleted using centrifugation and resuspended in fresh SM buffer.

Stained bacterial cells and stained phages were mixed together with a high MOI (~100) and directly analyzed using a Zeiss M2 imager fluorescence microscope.

### 2.14 Scanning electron microscopy

For scanning electron microscopy (SEM), bacteria were fixed with 3% (vol/vol) glutaraldehyde (Agar Scientific, Wetzlar, Germany) in PBS for at least 4 h, washed in 0.1 M Soerensen’s phosphate buffer (Merck, Darmstadt, Germany) for 15 min, and dehydrated by incubating consecutively in an ascending acetone series (30%, 50%, 70%, 90%, and 100%) for 10 min each and the last step thrice. The samples were critical point dried in liquid CO_2_ and then sputter coated (Sputter Coater EM SCD500; Leica, Wetzlar, Germany) with a 10-nm gold/palladium layer. Samples were analyzed using an environmental scanning electron microscope (ESEM XL 30 FEG, FEI, Philips, Eindhoven, Netherlands) with a 10-kV acceleration voltage in a high-vacuum environment.

### 2.15 Proteome analysis

#### 2.15.1 Sample preparation for proteome analyses

For the proteome analysis, propagated and CsCl-purified CL31 phages were used (described before). These phages were either pre-analyzed with SDS-PAGE [45] using a 4–20% Mini-PROTEAN® gradient gel (Bio Rad, Munich, Germany), or used directly for whole phage digestion.

#### 2.15.2 In-gel digestion

The de-staining and tryptic digestion of protein bands in gel slices was carried out in combination with the Trypsin Singles, Proteomics Grade Kit (Sigma-Aldrich) as previously described by Lavigne and colleagues [46]. The StageTipping desalting step was carried out as described elsewhere [47,48]. The tryptic peptide samples were stored at −20°C until use for MS measurements.

#### 2.15.3 Digestion of Whole Phage Particles

The digestion of whole phage particles (1 × 10^10^ PFU) were carried out as previously described by Lavigne and colleagues [46]. The StageTipping desalting step was carried out as described elsewhere [47,48]. The sample was dried and resuspended in 30 μL solvent A (H_2_O + 0.1% formic acid).

### 2.16 LC separation and mass spectrometry

#### 2.16.1 Shotgun proteomic measurement

The prepared tryptic peptide samples were separated chromatographically on a nanoLC Eksigent ekspert™ 425 LC system in microLC modus (Sciex) with a 25 Micron ESI Electrode coupled to a TripleTOF™ 6600 mass spectrometer (Sciex). As the trap, a YMC-Triart C18 capillary guard column with the dimension 5 x 0.5 mm ID, 12 nm, S-3 μm, was used. The analytical column was an YMC-Triart C18 column with 150 x 0.3 mm ID, 12 nm, S-3 μm (YMC). The column oven was set to 40 °C.

#### 2.16.2 Trapping conditions

The loading solvent was 2% acetonitrile in water with 0.5% formic acid. For the IDA (Information Dependent Acquisition) = DDA (Data dependent analysis) measurements, 8 uL or 10 uL of the desalted samples were loaded onto the trap column using 100% loading solvent for 3 minutes at 10 μL/min for desalting and enrichment.

#### 2.16.3 Separation conditions

The solvent used for mobile phase A was water with 0.1% formic acid and the mobile phase B was acetonitrile with 0.1% formic acid (both LC-MS grade, ROTISOLV®, ≥99.9%, Carl Roth). The separation of the peptides followed on the analytical column with a linear gradient with increasing concentration of mobile phase B. The initial conditions were 97% A and 3% B at a flow rate of 5 μL/min. The linear gradient used was 3% - 25% B in 38 minutes. Within the following 5 minutes, the mobile phase B was increased to 32% and was further increased to 80% within 2 min, lasting for 3 minutes. At minute 49, the mobile phases were changed to the initial condition with 97% A and 3% B within 8 min.

#### 2.16.4 IDA-MS

The source and gas settings were 5500 V Ionspray voltage, 35 psi curtain gas, 12 psi ion source gas 1, 20 psi ion source gas 2 and 150°C interface heater.

For shotgun measurements the mass spectrometer was operated in positive mode with a “top 50” method: Initially, a 250 ms survey scan (TOF-MS mass range m/z 400-1250), was collected from which the top 50 precursor ions were automatically selected for fragmentation, whereby each MS/MS event (mass range m/z 100-1500, in high sensitivity mode) consisted of a 50 ms fragment ion scan. The main selection criterion for parent ion selection was precursor ion intensity. Ions with an intensity exceeding 150 counts per second and with a charge state of +2 to +5 were preferentially selected. Those precursors were added to a dynamic exclusion list for 15 s. Precursor ions were isolated using a quadrupole resolution of 0.7 amu and fragmented in the collision cell with a rolling collision energy of 10 eV. If less than 50 precursor ions fulfilling the selection criteria were detected per survey scan, the detected precursors were subjected to extended MS/MS accumulation time to maintain a constant total cycle time of 2.8 s.

#### 2.16.5 Data analysis

The IDA data were processed with ProteinPilot™ (5.02, Sciex) using the Paragon™ algorithm for protein identification This data was then compared with a database using a protein FASTA file of *C. glutamicum* ATCC 13032 and predicted protein sequences of the phage CL31.

## 3. Results

### 3.1 Genome assembly, annotation and comparisons

The corynephage CL31 was already characterized with regard to its structure and restriction map in 1987 by Annie Trautwetter and colleagues [21]. In this study, we set out to further characterize this Siphovirus and its interaction with the biotechnological platform strain *C. glutamicum* ATCC 13032.

Genome sequencing and assembly revealed that CL31 harbors a 45,061 bp dsDNA genome with an average GC-content of 54.4% (Figure 1A). Using the bioinformatics tool PhageTerm, the termini of CL31 were identified as being cohesive ends with a 16 bp 3’ overhang (CCCCACTCGGCCCACG)[37]. Genome annotation revealed 72 genes encoded by the CL31 genome (Figure 1A and Table S3). While nearly half of the discovered genes encode proteins of unknown function (34/72), a putative function could be assigned to the rest of the CL31 proteins(Figure 1B). A main group consists of genes encoding phage structural proteins (clg52 – clg72). Together with phage lysis genes (clg49 – clg51), this group represents a major proportion of all coding sequences on the CL31 genome (Figure 1A). Besides these phage-related genes, CL31 also encodes two “diverse” sets (clg01 – clg28 and clg29 – clg48) of genes that mainly consist of genes related to DNA replication, repair, recombination and degradation as well as with amino acid transport and metabolism. While the first diverse set is solely encoded on the minus strand, the second set is encoded on the plus strand of CL31.

**Figure 1:**
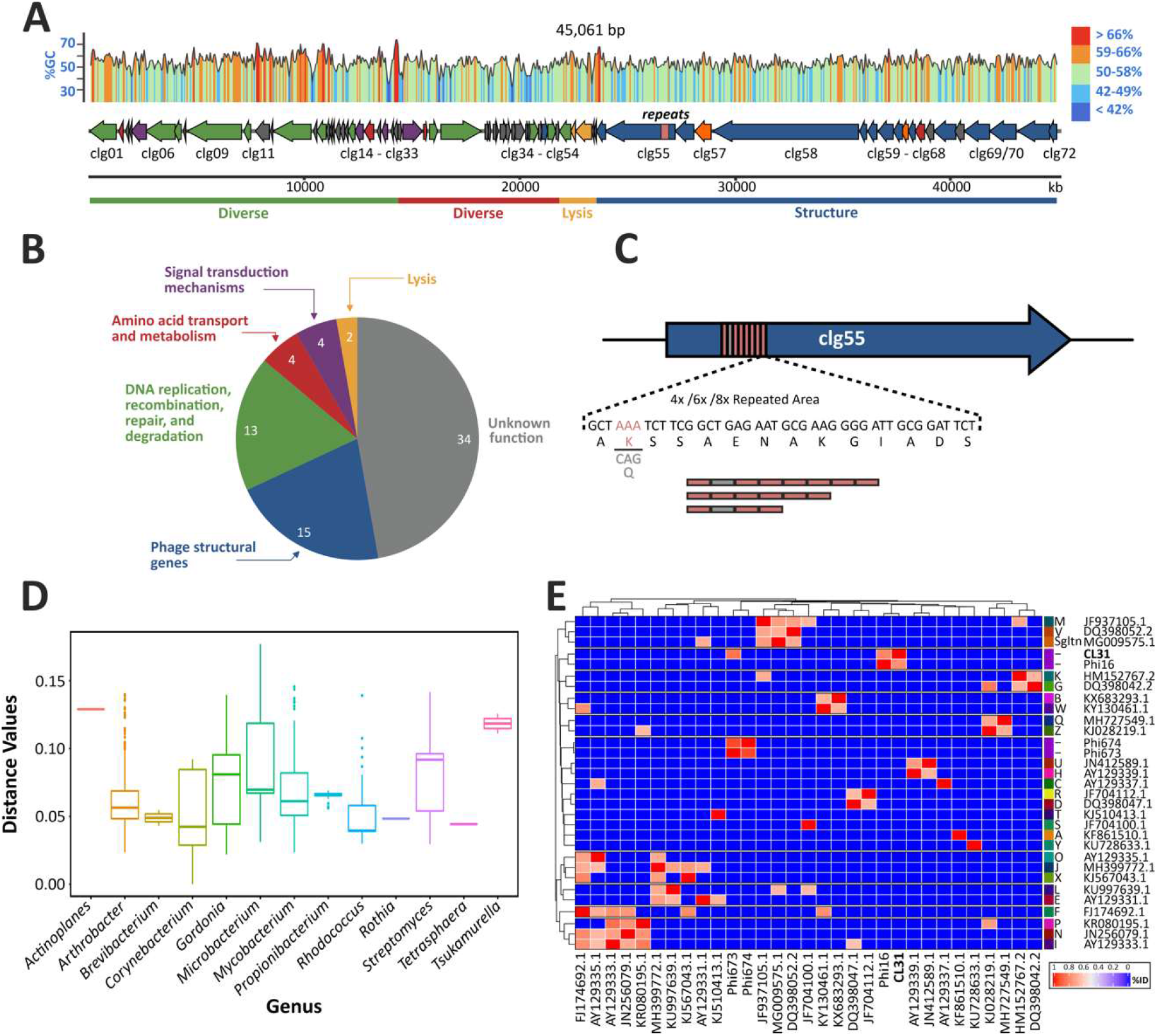
Genome analysis of the corynephage CL31. (A) The genome of CL31 was assembled based on a combination of Illumina sequencing and Sanger sequencing. ORF prediction was done using Phanotate, GeneMark and SnapGene. The annotation and functional categorization (B) are based on multiphate as well as BLASTp against nr database and the PhagesDB database (phagesdb.org). (C) The gene clg55 contains a variable repeat region consisting of 42 base pair minisatellites. This region could not be assembled using formerly described assembly methods but was resolved by Sanger sequencing. (D) Pairwise distance between CL31 and >2950 phage genomes based on the Jensen-Shannon divergence method (E) Heatmap representing an average nucleotide percentage identity matrix with distances obtained using PYANI with the BLAST (ANIb) method based on a selected set of 31 phage genomes, including CL31.

Interestingly, during genome assembly an area was discovered showing uncertain read mapping properties and reads that only partly mapped inside the gene clg55 encoding a putative minor tail protein (data not shown). To discover the origin of this mapping conflict, we used a primer pair flanking the particular genomic region to perform PCRs on single plaques and Sanger sequencing to resolve the correct sequence of this area (Figure S1). Remarkably, different plaques led to different sizes of the PCR products suggesting that this area could be variable within the phage population. In line with this observation, Sanger sequencing revealed the occurrence of three different lengths of a repeat area inside the gene clg55 (Figure 1 C and Figure S1). The three exemplary plaques picked contained - as schematized in Figure 1C - either four, six or eight repeats inside of clg55. These repetitive areas consist of single elements of 42 bp minisatellites, spanning 14 amino acids. In case of the four or eight repeat variants, the second microsatellite differs from the others because of a codon exchange (AAA to CAG), shown in Figure 1 C in light grey.

Pairwise comparison of the CL31 genome against 2955 previously described different actinophage genomes was performed to describe the distance between CL31 and other phages (Figure 1D). This analysis confirmed that CL31 features the highest similarity to other *Corynebacterium* phages followed by phages infecting species of the genus *Gordonia*, *Mycobacterium* and *Arthrobacter* (Figure 1D and Table S4). An ANI-based distance matrix (Figure 1E) showed that the most similar phage to CL31 is the temperate corynephage Φ16 [9,11]. Another corynephage showing low similarity to CL31 is the lytic phage Φ673 [14], but no clustering was detectable with other exemplary phages of each cluster (clustering according to phagesDB, based on [49]).

### 3.2 Plaque morphologies and infection dynamics of CL31

In the following, we investigated the plaque morphology and infection dynamics of CL31 on different *C. glutamicum strains*. A first set of experiments revealed that CL31 showed the most efficient propagation when infecting the prophage-free *C. glutamicum* ATCC 13032 derivative MB001 (designated as MB001 in the following). In comparison, the WT strain ATCC 13032 lead to an EOP of 1.7 10-6 and the same strain lacking the restriction modification system CglIM-CglIR-CglIIR (Δ*resmod*) to an EOP of 0.63. A magnified view on plaques appearing on a MB001 lawn with CL31 revealed the occurrence of two distinct types of plaques: on the one hand slightly bigger and turbid ones, and on the other hand smaller plaques with a clear center and rougher border (Figure 2A). While CL31 was predicted to be a temperate phage (Table S6), the appearance of clear plaques could hint on the generation of lytic mutants that lost the ability to form lysogens with the host strain.

**Figure 2:**
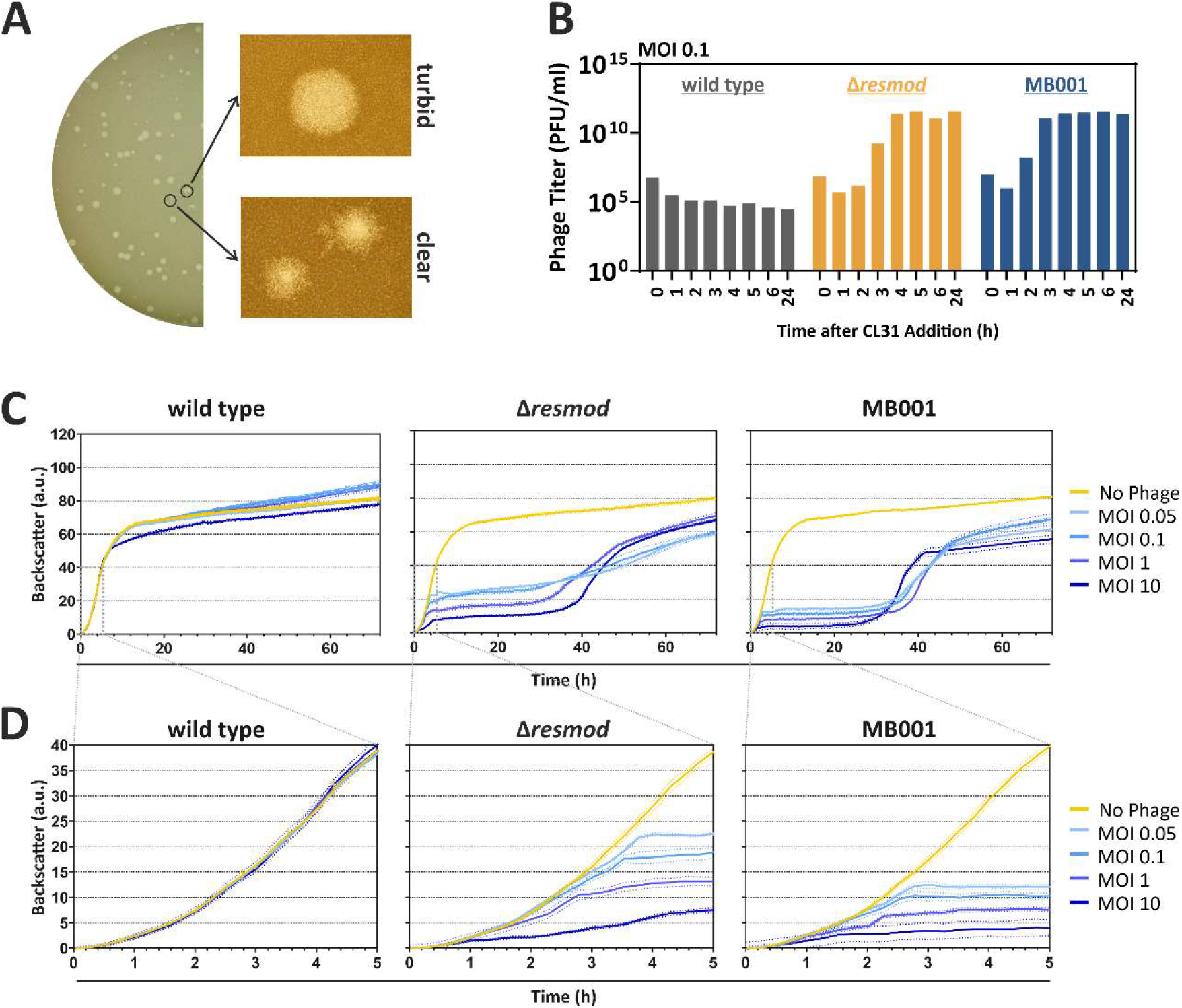
Plaque morphology and infection dynamics of CL31. (A) MB001 cells (OD_600_ 0.5) were mixed with 10 μl CL31 phages (stock diluted to 10-9) in an overlay agar. After over-night incubation, plates were imaged using a Nikon SM-18 stereomicroscope. (B) Infection assay of CL31 with different host strains was used to determine the phage titer at different time points of incubation. For this purpose, bacterial cells were incubated with CL31 in BHI medium applying an initial MOI of 0.1. At the described time points, samples were taken and 3 μl of the centrifuged supernatant were spotted on a MB001 loan (OD_600_ 0.5); plaques were quantified after overnight incubation (see also Figure S2). (C) Growth of different *C. glutamicum* host strains infected with different MOIs of CL31 using a BioLector microcultivation system. All data represent mean values with standard deviations from three independent biological triplicates (n=3). (D) Zoom into the first 5 hours of cultivation shown in (C).

Infection assays in liquid cultures were performed in shaking flasks to monitor the amplification of phage titer (Figure 2B) as well as in a Biolector microcultivation system (Figure 2C and 2D). For all tested strains, an initial decrease in phage titer is visible up to 1 h after start of the cultivation. Because the titer determination is based on free phages in the medium, this decrease probably reflects the adsorption of the free phage virions to the host cells. In this assay, no significant amplification of CL31 was observed for the wild type strain. In contrast to this, the Δ*resmod* strain as well as the prophage-free strain MB001 showed an increase in phage titer starting already after 2 h. Independently of the starting MOI, both strains reached a maximum titer of 10^12^ PFU/ml (Figure 2B and Figure S2). Remarkably, phage amplification was clearly delayed in *C. glutamicum* Δ*resmod* in comparison to strain MB001 (maximum in titer after 4 versus 3 hours, respectively). This altered infection dynamics is also reflected by the growth of the different strains in the presence and absence of phages: While *C. glutamicum* wild type was only slightly affected in growth with the highest MOI of 10, strains Δ*resmod* and MB001 already displayed strong growth defects at an MOI of 0.05. Analogously to the delayed increase of phage titer in shake flask experiments of observed for *C. glutamicum* Δ*resmod*, growth of strain MB001 plateaued significantly earlier (Figure 2C and 2D). Independently of the used MOI, both sensitive strains (Δ*resmod* and MB001) showed resumed growth after approximately 30 hours after infection probably hinting at the presence and growth of CL31-resistant clones or the formation of lysogens (Figure 2C).

### 3.3 Visualization of CL31 attachment and lysis

Fluorescence microscopy (Figure 3A) and scanning electron microscopy experiments (Figure 3B) were used to visualize attachment and CL31-induced lysis of strain *C. glutamicum* MB001. To this means, phage particles were stained using the nucleic-acid stain SYBR Gold (Figure 3A). In addition, host cell membranes were stained using Nile red; the nucleoid was visualized using Hoechst 33342. After adding phages to a fresh culture of host cells, pictures were taken with a fluorescence microscope. Figure 3A clearly shows the occurrence of green foci on the bacterial surface, which hints on the attachment of stained CL31 phages to the bacteria. Free phages were also detectable in the surroundings of the bacterial cells. In addition to the surface-attached signals and the free phage particles, a green staining of the host chromosome was detectable. This signal, however, likely results from a carryover of the SYBR Gold stain, since it was detected immediately after addition of the phage sample.

**Figure 3:**
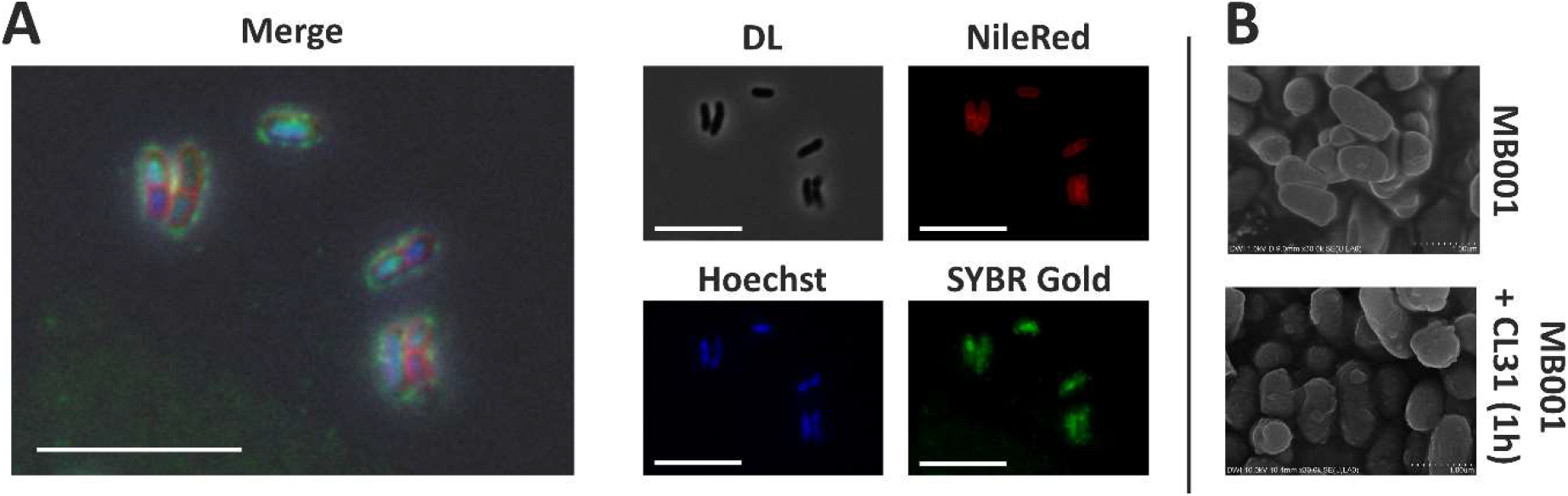
Visualization of CL31 attachment and host lysis. (A) Fluorescence microscopy of *C. glutamicum* MB001 cells stained with Nile red and Hoechst 33342, mixed with SYBR Gold stained CL31 phages. The bacterial cells and phages were mixed with a high MOI (~100) and directly analyzed using a Zeiss M2 imager fluorescence microscope. (B) Scanning electron microscopy images taken from *C. glutamicum* MB001 cells without (top) and with (bottom) CL31 phages. Bacterial cells and phages were mixed with an MOI of 1.2 and incubated for 1 h at 30°C and 120 rpm in BHI medium.

The scanning electron microscopy images (Figure 3B) was further used to visualize phage-induced cell lysis. After one hour, cells infected with CL31 showed a disintegrated cell morphology and bubbles on the cell surface indicating the start of cellular lysis.

### 3.4 Global transcriptome of C. glutamicum during infection with CL31

To assess global changes on the level of the transcriptome, we conducted an RNA-seq analysis with infected and uninfected *C. glutamicum* wild type and MB001 (Figure 4 and Figure S3). Successful infection was confirmed by monitoring growth and plaque-forming units (Figure S3A). To cover the time point of highest intracellular phage activity, we decided to analyze the transcriptome of the samples taken 1 h after infection. At this time point, the amount of extracellular phage particles was the lowest before the titer increased again after 1.5 hours (Figure S3B). A comparison of the read origin of all tested samples shows the distribution of all reads to either host genes, genes inside of the CGP3 prophage or CL31 genes (Figure 4A). As both of the tested strains contain a similar fraction of host genes, the distribution is equal in this case. For CGP3, however, only the wild type strain shows a high signal, because this prophage region was deleted in MB001. In line with the above-described experiments, CL31 transcripts could only be detected for MB001.

**Figure 4:**
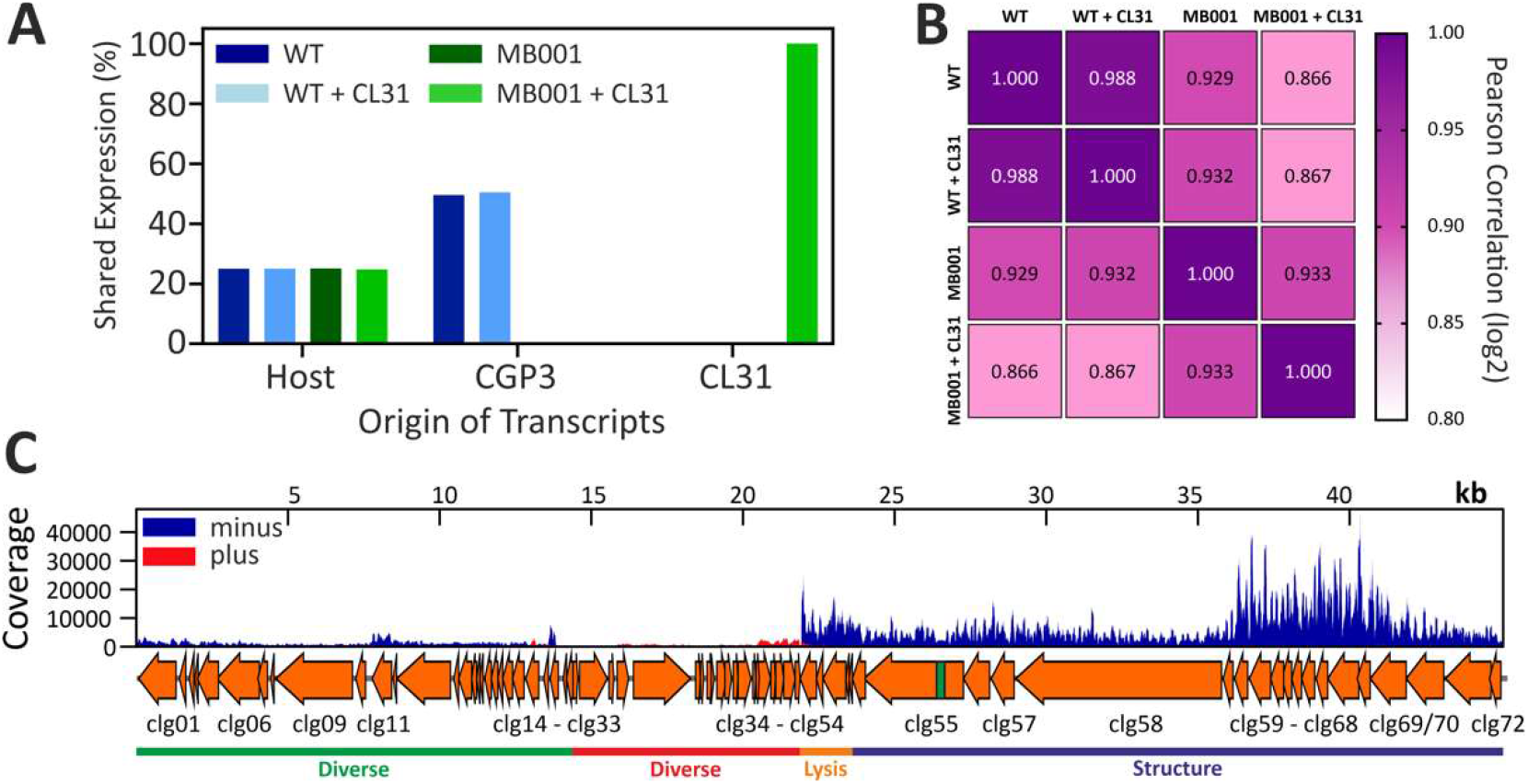
Transcriptome analysis of CL31-infected *C. glutamicum* strains. (A) The origin of transcripts is displayed as relative part of each sample. It was divided into transcripts from the host, from the CGP3 prophage element of the host or into transcripts from the CL31 phage. Bars represent mean values of two replicates. (B) Pearson correlation was calculated based on the Log2 of the TPM values of each experiment. (C) Transcription profile of the CL31 phage infecting *C. glutamicum* MB001. Bacterial cells and phages were mixed with an MOI of 0.1 and incubated in BHI medium at 30°C and 120 rpm. The coverage shows one representative sample. A detailed sampling is shown in Figure S3 and detailed analysis is represented by Table S5.

This effect is further underlined by a comparison of the transcriptional correlation of both infections (Figure 4B). While no strong differences are detectable in case of wild type with and without CL31 infection, the transcriptome of *C. glutamicum* MB001 differed significantly when infected and uninfected samples were compared (Figure 4B). Furthermore, a strain-dependent difference is detectable between uninfected wild type and MB001 cells.

The analysis of CL31-derived transcripts (Figure 4C) showed a strong transcription of genes located on the minus strand (especially clg49-clg72) but only slight signals from the plus strand (clg24-clg48). The majority of CL31 genes transcribed at 1 h after infection mapped to genes encoding structural components of the viral particles as well as lysis proteins (Figure 4C and Table S5). This suggests a late infection state where new phage particles are already assembled and expression of clg51 (lysin) and clg50 (holin) will initiate cell lysis. Besides structural and lysis genes, the highest transcription was observed for a Xre-/Cro-like repressor gene (clg26). However, all genes surrounding clg26 are either not transcribed or only at lower levels. Table S5 shows further that only 8/72 genes do not show any transcription at this time point. This covers the genomic regions from clg27 to clg39 where genes are mainly located on the plus strand of CL31. Except a potential antirepressor protein (clg30) and a peptidase (clg31), all of these non-transcribed genes code for hypothetical proteins (cp. Table S3).

Taking together, the transcriptome analysis revealed a successful infection of MB001 but not of the wild type, which is in agreement with the infection curves showing no significant amplification of CL31 in *C. glutamicum* WT cells.

In addition to the CL31-derived transcripts, aim of our analyses was the host reaction towards infection with this phage. Table S5 contains – in addition to the CL31 transcriptome - the whole transcriptomes of both MB001 and the WT strain. As a reaction to the CL31 addition, the expression of 156 genes in total was more than 2-fold up or down regulated (Table S5), while in MB001 expression of 122 genes are affected.

In the case of *C. glutamicum* WT infection, genes affected by CL31 infection belong to diverse functional groups including genes involved in transport and metabolism (carbon sources, ions, and metabolites), DNA replication and metabolism, or signal transduction mechanisms. Interestingly, 23% of the upregulated and 18% of the downregulated genes are located within the CGP3 region. The highest upregulated gene of the CGP3 region is cg1988 (5.6-fold upregulated) encoding a putative transcriptional regulator with similarities to a phage immunity repressor protein. Additionally, the neighbouring gene cg1989, which shows similarities to a metallopeptidase, is also upregulated during CL31 infection (cp. Table S5).

### 3.5 Structural proteome analysis of purified CL31 virions

A significant fraction of CL31 CDSs on the minus strand encodes proteins potentially serving as structural components of the CL31 virion. To confirm the presence of these protein in the phage particle, CL31 virions were purified using CsCl-gradient centrifugation. Samples were analysed by SDS-PAGE as well as mass-spectrometry analysis of denatured phage particles (Figure 5). The combined results from both approaches are depicted in Figure 5C.

**Figure 5:**
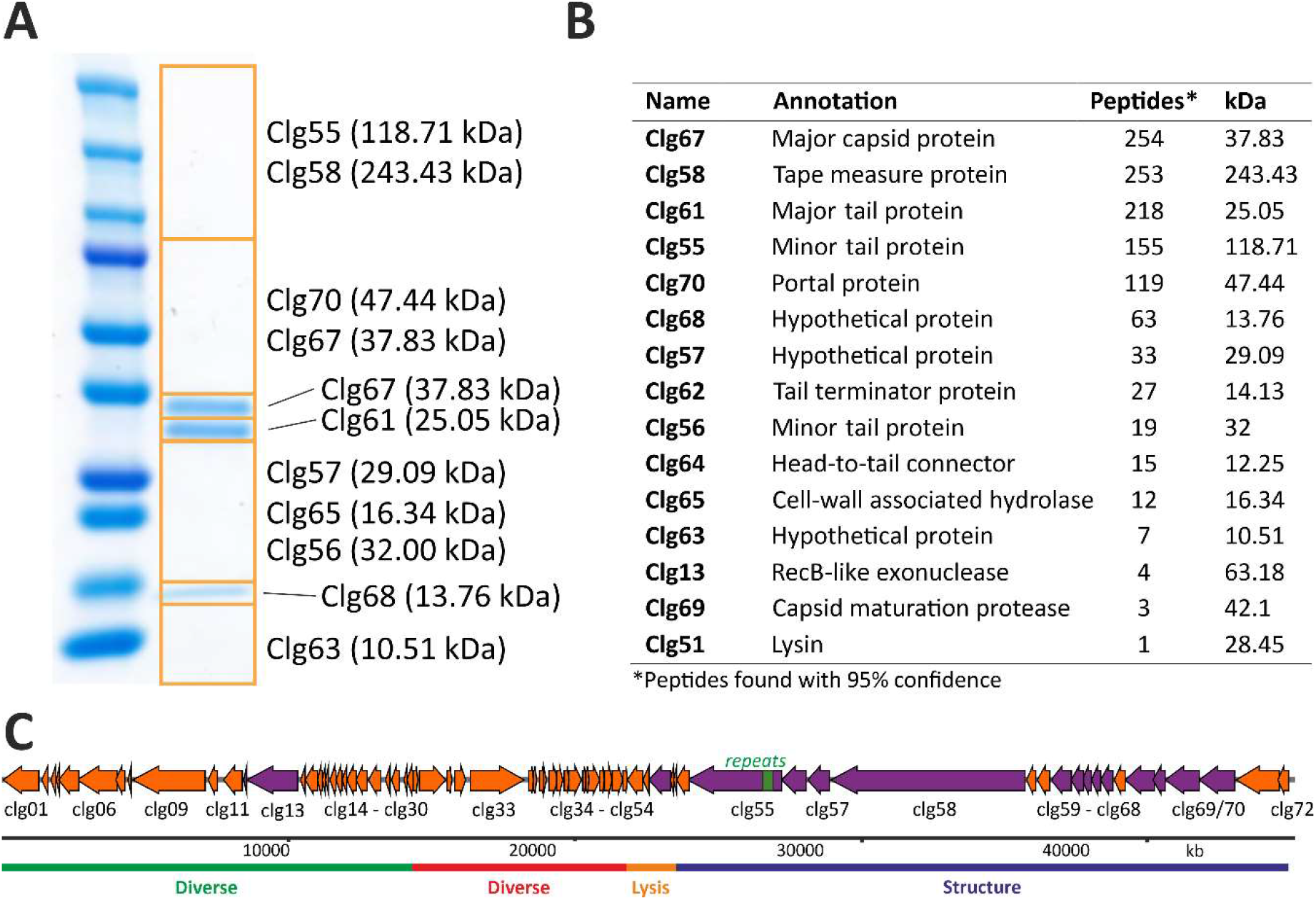
Analysis of the structural proteins of CL31 using SDS-PAGE and mass spectrometry. (A) SDS-PAGE of CL31 phage particles purified with CsCl-gradient centrifugation. (B) LC/MS analysis of whole phage particles. (C) Comprehensive scheme of all proteins found (purple) in (A) and (B).

As expected, the majority of detected proteins could be attributed to the group of structural proteins. Two exceptions are represented by Clg13 (RecB-like exonuclease) and Clg51 (lysin) (Figure 5C). However, these proteins were found in minor fractions and might be remnants of the cell lysis. Overall, the identified structural protein components nicely correspond to the composition of a full phage particle. The phage tail structure is likely composed of the minor tail proteins Clg55 and Clg56, the tape measure protein Clg58 and the major tail protein Clg61, while the phage capsid consists of the major capsid protein Clg67. Major tail and major capsid proteins (Clg67 and Clg61) were also detected as clear bands on the SDS-PAGE (Figure 5A). Additionally, several further phage structure proteins could be identified at lower abundances. These included a tail terminator protein (Clg62), a head-to-tail connector protein (Clg64), a capsid maturation protease (Clg69) and a portal protein (Clg70). Other proteins found and encoded in the structural protein part of CL31 are Clg65 (put. cell-wall associated hydrolase) and several hypothetical proteins of unknown function.

### 3.6 Mutations in genes involved in mycolic acid biosynthesis lead to resistance to infection by CL31

A prolonged incubation of *C. glutamicum* host cells gave rise to a cell population able to grow in the presence of phages after ~30 h. Interestingly, isolation of clones surviving phage infection showed an altered colony morphology after the cultivation (Figure 6A). While WT cells showed a homogeneous and smooth colony surface structure, survivors of CL31 infection displayed a rough and uneven colony surface. Resistance of isolated clones obtained from all three *C. glutamicum* strains (wild type, Δ*resmod* and MB001) was first verified using spot assays (Figure 6B). To identify causal mutations conferring resistance to phage infection, we performed genome sequencing of single phage-resistant clones. Genome sequencing of all three clones was conducted, followed by an identification of single nucleotide polymorphisms (SNPs) using the bioinformatics tool Snippy [50]. As shown in Table 2, in comparison to the parental strains (*C. glutamicum* ATCC 13032, accession: BX927147 or *C. glutamicum* MB001, accession: CP005959), each of the survivors showed only one SNP. All SNPs caused missense mutations leading to an amino acid exchange of either glycine or tyrosine to aspartic acid (Table 2) and were found in genes relevant for the biosynthesis of mycolic acid in *C. glutamicum*. Two mutations were found in *accD*2 and *accD*3 encoding subunits of a acyl-CoA carboxylase complex [51]. The third mutation was identified in *pks*, encoding a polyketide synthase. All three gene products were previously described to share important functions in precursor supply for mycolic acid biosynthesis [51]. As a verification of the resistance phenotype caused by the mutations, all found mutations were re-integrated in the most sensitive strain – MB001. Spot assays of CL31 on lawns of the isolated mutant strains as well as strains carrying the re-integrated single SNPs confirmed an almost complete resistance of the respective strains towards CL31 infection (Figure 6C).

**Table 2:**
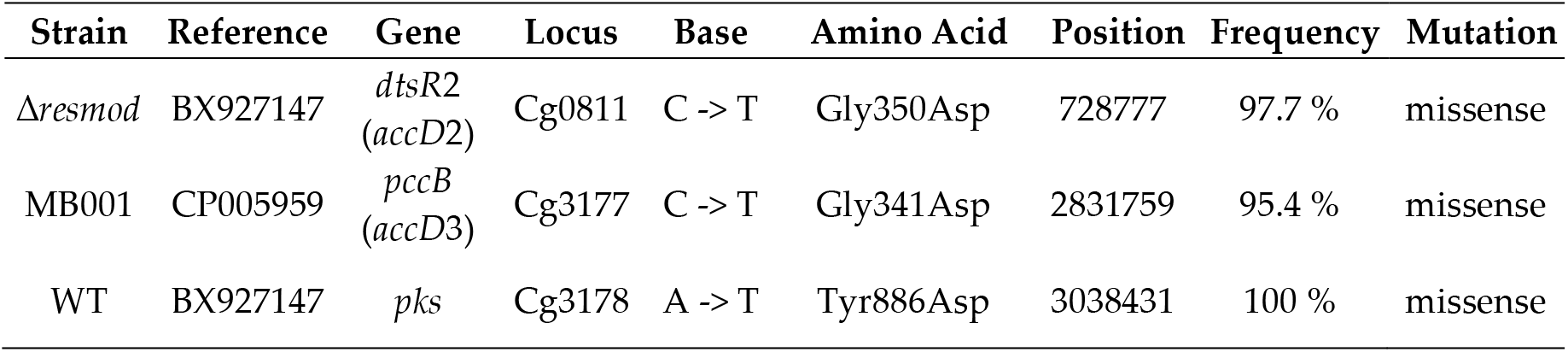
SNP analysis using the bioinformatic tool Snippy [50] was used to discover mutations in different *C. glutamicum* strains showing resistance towards CL31 infection.

**Figure 6:**
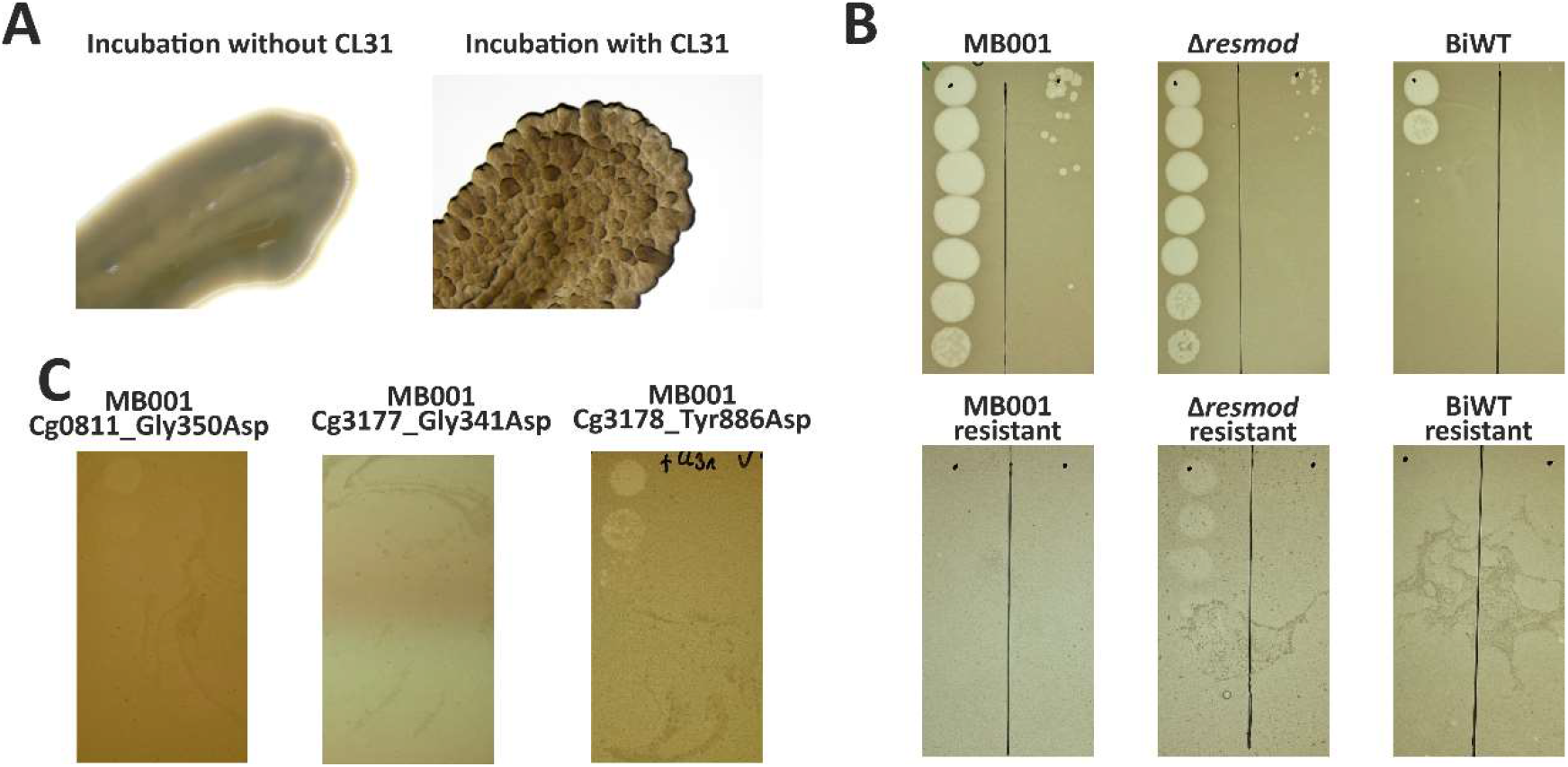
SNPs in genes involved in mycolic acid biosynthesis confer resistance to CL31 infection. (A) Colony surface of C. glutamicum WT (left) and a clone isolated after CL31 infection (right). For details on the isolation, see material and methods. (B) *C. glutamicum* clones isolated after CL31 were used to inoculate an overlay agar and perform phage titer determination with a serial dilution of CL31. As controls, the corresponding origin strains were treated similarly (top row). Representative examples out of three replicates are shown. (C) A similar titer determination as described in B was conducted with *C. glutamicum* strains harboring a reintegration of the SNPs (Table 1) found in the resistant *C. glutamicum* strains.

## 4. Discussion

The present study provides a comprehensive analysis of the genomic features of phage CL31 and the infection of the host strain *C. glutamicum* ATCC 13032. While the infection of the WT strain ATCC 13032 was only poorly visible with spot assays and no phage amplification occurred, adsorption of CL31 to the WT was still detectable (Figure 2). In contrast, *C. glutamicum* Δ*resmod*, lacking the CGP3-encoded restriction modification system, showed a clear propagation of CL31, thus, confirming the central role of the CGP3-encoded restriction modification system in phage defense [52]. Interestingly, the strain MB001 lacking all prophage elements, revealed a significantly accelerated phage amplification suggesting that so far, unknown elements of the cryptic prophages might collude with their host to confer resistance to phage infection.

The additional effect conferred by the presence of prophages can be in principle explained in two ways; (i) a prophage-encoded protein or RNA is interfering with CL31 amplification or (ii) the induction of the CGP3 prophage in a fraction of *C. glutamicum* cells is limiting the spread of the infection [53]. While the latter ‘abortive infection’-like mechanism was also hypothesized as important mechanism for different *C. glutamicum* strains to resist infection of BFK20 [54], examples of prophages assisting their host to resist superinfection have been described for different mycobacteriophages [55,56]. Besides specific restriction modification systems, Dendrick and colleagues found prophages encoding cell envelope modifying systems as well as prophages causing repressor-mediated immunity. Furthermore, special and novel defense systems were found encoded on prophages. One example is the Gp30/Gp31 system of the temperate mycobacteriophage Sbash [57]. Gentile and colleagues demonstrated the ability of this system to interrupt lytic growth of mycobacteriophage Crossroads using an ion channel to destroy the membrane potential and thus leading to a reduction of ATP synthesis.

One interesting target with a potential role in the recognition of foreign DNA, which is encoded on the CGP3 element, is the xenogeneic silencer CgpS. In previous studies, CgpS was described as a small nucleoid-associated protein binding to multiple targets within the cryptic prophage elements of *C. glutamicum* [5,58]. However, tests with MB001 containing this silencing protein did not show any difference in infection dynamics compared to a strain carrying an integrated copy of *cgpS* (data not shown). An explanation for this observation probably lies in the binding properties of CgpS. As shown by Wiechert and colleagues, binding of CgpS requires high AT content as well as a specific binding motif [59]. CL31, in contrast, shows a similar GC content as the host (54 %) and thus might be able to escape the silencing by CgpS. Nevertheless, transcriptome analysis of WT cells infected with CL31 revealed some changed transcription of different prophage targets. Especially interesting could be the gene cg1988 encoding an immunity repressor and its neighboring gene cg1989 encoding a protein with similarities to a metallopeptidase. Pairs of immunity repressor and metalloprotease were recently shown to act as regulatory elements for prophage elements in *Listeria monocytogenes* [60]. Here, the metalloprotease encoded by a bacteriocin locus was shown to be upregulated in response to stress and acts as an anti-repressor protein to overcome the repression caused by a CI-like immunity repressor [60].

Another reason for the fast amplification of CL31 in strain MB001 could be the absence of lysogen formation. An integration of temperate phages into the host chromosome requires different essential factors, i.a. an integrase protein [61]. Because the CL31 genome does not encode an integrase protein on its own, this phage might be able to use a host-encoded integrase. The WT strain and the Δ*resmod* strain both contain the CGP3 prophage element encoding two putative integrase subunits [3]. This CGP3 prophage element was shown to be able to excise from the host genome underlining the presence of a functional integrase. Thus, the difference in sensitivity of MB001 and Δ*resmod* might also be caused by the formation of a small amount of lysogens that consequently lead to a super-infection exclusion of these lysogens [62].

Although this study could not experimentally prove the presence of CL31 lysogens, there are multiple aspects hinting at a temperate lifestyle of CL31. Among those is the fact that CL31 typically shows a turbid plaque morphology. In some cases, small and clear plaques occur suggesting the formation of lytic CL31 mutants [63]. Furthermore, a comparison of CL31 with other *C. glutamicum* infecting phages showed the highest similarity to the temperate phage Φ16 (Figure 1).

Sequencing of strains resistant to CL31 infection, interestingly, showed SNPs converging in pathways required for the synthesis of mycolic acids. Mycolic acids are a central component of the cell envelope composition of *C. glutamicum*. Here, trehalose monocorynomycolate (TMCM) and trehalose dicorynomycolate (TDCM) represent the most abundant lipids in the *C. glutamicum* ATCC 13032 [64]. Together with arabinogalactan-bound mycolates they form the outer layer of the *C. glutamicum* envelop, which is part of the mycoloyl-arabinogalactan-peptidoglycan (mAGP) mesh [65]. Houssin et al. further described this outer membrane as spanned by e.g., porins and partly modified with an outer layer mainly consisting of glucan. For the biosynthesis of mycolic acids, two acyl-CoA carboxylase subunits (AccD2 and AccD3) as well as a polyketide synthase (*Cg*-pks) are essential key enzymes [51,66]. Fatty acid precursors are activated by either fatty acyl AMP ligase (FadD) or acyl-CoA carboxylase complex (AccBCD2D3E) and further processed by Cg-Pks that catalyzes their condensation by a Claisen reaction, forming an α-alkyl β-ketoester which is further reduced to a mycolic acid [51,67]. Astonishingly, different CL31-resistant strains carried SNPs in each of the three genes (*accD*2, *accD*3, and *Cg-pks*). All mutations were missense mutations leading to the substitution of a glycine (*accD*2, *accD*3) or a tyrosine (Cg-pks) with aspartic acid. Because of the structural differences between the original amino acid and the substitution, it can be assumed that the enzymes are impaired in their function. Similar to our observation for CL31-resistant strains, Gande and colleagues also described a rough colony surface of strains lacking one of the indicated genes (*accD*2, *accD*3, and *Cg-pks*) [51]. Additionally, they reported a loss of extractable mycolic acids and cell wall-bound mycolic acids. A change in mycolic acid composition was previously shown to cause a phage resistance phenotype in *Mycobacterium smegmatis* [68]. Based on these data, it is, however, not yet clear whether mycolic acids are directly recognized by phages as a first adhesion layer or if the loss of receptor proteins, such as porins or transporters embedded in the outer layer of the cell envelope is responsible for the resistance.

Future studies along these lines will target the molecular mechanism of host cell recognition by CL31 and the escaping strategies of the host. The latter also spans the aim of elucidating the consequences of the molecular arms race between phage and bacterium with regard to envelope modifications and bacterial resistance towards environmental stresses (as e.g., described for antibiotic vulnerability of *M. tuberculosis* when expressing a certain SWU1 phage gene [69]). A loss of the external mycolic acid layer might prevent bacterial cells from phage infection, but at the same time enhance sensitivity to certain antibiotic treatments [70].

One further interesting observation is the differences in the tandem repeat region inside the putative minor tail protein Clg55 (Figure 1). As described before, corynomycolates can have different lengths depending on the growth condition [71]. If these mycolic acids are crucial for phage recognition, the repeat region inside the minor tail protein might represent an adaptation to different host mycolic acid compositions. Interestingly, de Jong and colleagues could identify tandem repeats containing a BACON (*Bacteroides*-associated carbohydrate-binding often N-terminal) domain inside different phage tail proteins of crAss-like phage lineages infecting *Bacterioides* species. They hypothesize a possible function in host recognition [72]. The architecture of the BACON domain was associated by Mello and colleagues as being involved in carbohydrate binding [73]. Remarkably, a domain prediction using the online tool InterPro [74] revealed that Clg55 also contains a galactose-binding superfamily domain (data not shown). In a recent study focusing on *Brevibacterium aurantiacum* phages, De Melo and colleagues detected tandem repeat areas in 85% of all phage genomes available in public databases [75].

In summary, this work provided comprehensive insights into the genomic features of phage CL31 and its interaction with the biotechnological platform organism *C. glutamicum* ATCC 13032. Further mechanistic studies will aim at the development of novel tools for metabolic engineering and synthetic biology applications.

## Acknowledgments

We thank Osher Fiyaksel from Prof. Sigal Ben-Yehuda’s lab at the Hebrew University of Jerusalem, Israel, for assistance with the phage staining protocol. Furthermore, we thank Andrei Filipchyk for his help with statistics and bioinformatic analyses. Our thanks also go to the entire Frunzke lab for constant friendly support and all stimulating discussions.

