## supplemental Information for "Genome sequencing of the bacteriophage CL31 and interaction with the host strain *Corynebacterium glutamicum* ATCC 13032"

Genome sequencing of the bacteriophage CL31 and analysis of its characteristics and host interactions

Received: date; Accepted: date; Published: date

List of all supplementary files

Table S1: Oligonucleotides used in this study.

Table S2: Construction of plasmids used in this study.

Table S3 (excel): Gene prediction and annotation of the CL31 genome.

Table S4 (excel): Distances of CL31 in comparison with other phage genomes.

Table S5 (excel): Transcriptome analysis of host cells and CL31 during infection of MB001 cells.

Table S6: Lifestyle prediction and validation of CL31 and other corynephages.

Figure S1: PCR and Sanger sequencing of singles plaques to close the CL31 genome.

Figure S2: Spot assays of the infection dynamics analysis of CL31 with different hosts.

Figure S3: Infection curve for RNA-seq analysis.

**Table S1.** Oligonucleotides used in this study.

| **Number** | **Sequence (5’-3’)** |
| --- | --- |
| 1 | CCTGCAGGTCGACTCTAGAGAGCATGGAATCGCCACCGAT |
| *2* | CATTCCACTATAGAAACTGTTTTTTAAAGGAGAACCATGTCTGA |
| *3* | TAAAAAACAGTTTCTATAGTGGAATGTTACCGTGCTTTTTAGC |
| *4* | CCCAGTTCGCTGACTGCTTGGATAT |
| *5* | CAAGCAGTCAGCGAACTGGGTTGGC |
| *6* | CTAAGAAACCATGACTGCAGCACAGACCA |
| *7* | CTGCAGTCATGGTTTCTTAGTCTAAATCCTGAAGG |
| 8 | TTGTAAAACGACGGCCAGTGAGACCACATGCTCAACCCGT |
| 9 | CCTGCAGGTCGACTCTAGAGgaggtctcgcacagccttggt |
| 10 | gccgacaaggacgcccgcttcatcc |
| 11 | aagcgggcgtccttgtcggcggcgt |
| 12 | TTGTAAAACGACGGCCAGTGacctgtgaccactgactttgttatcggcgt |
| 13 | CCTGCAGGTCGACTCTAGAGAATCTGAGGGCTGCGGCC |
| 14 | GGAAGAATTAGTAACGGAGAGCTGACGGAAG |
| 15 | CTCTCCGTTACTAATTCTTCCGAGAAATCTCATCAATGC |
| 16 | AGTGGTCCACGACGCCGATGACATG |
| 17 | CATCGGCGTCGTGGACCACTGCTCC |
| 18 | AGGCCTCAATATGGAACAGAGCCAAT |
| 19 | TCTGTTCCATATTGAGGCCTCAGCCT |
| 20 | TTGTAAAACGACGGCCAGTGCGCTCCGCCGTTGGC |
| 21 | GTGTCCAATTCCAGATCCACATCGT |
| 22 | ACGATGTGGATCTGGAATTGGACAC |
| 23 | CTGGTTGGTGCCAGGCAGGAAG |
| 24 | GGTGGTCACGTGTACTCCCCTG |
| 25 | CTGCAGCCTTGGTGTCCTGAGT |
| 26 | CACACAAGAACCCTGCACAACGC |
| 27 | GGTCATGCATGTCGTAGCCCTGGTTA |
| 28 | TAACCAGGGCTACGACATGCATGACC |
| 29 | CCACGACGAAGATGATCGGGAT |
| 30 | ATGTATGTCACCGGCCCAGCC |
| 31 | CCCGACAGAACTGCGAAGATGG |
| 32 | GGAGCTCATTTGAGTCGCCGC |
| 33 | GCATGTTCTTCTTCTCTGGTAGCGG |
| 34 | CGCGATACCATGCTCATTGCCAG |
| 35 | GTCGCTGACGTGGTAATCATGGTGG |
| 36 | CGGATCCTTTGTGGAGATCGGCG |
| 37 | CCGCTACCAGAGAAGAAGAACATGC |
| 38 | CTGGCAATGAGCATGGTATCGCG |
| 39 | CCACCATGATTACCACGTCAGCGAC |
| 40 | CGCCGATCTCCACAAAGGATCCG |
| 41 | GATTTCAATTCCGGAACGCTCGGTGAG |
| 42 | GGCTACGACGGATTCAACACCGAT |
| 43 | ATGCCAGCGATTTCGCCAGC |
| 44 | CGAGGAAGCCGTAAAGCGCC |
| 45 | GCGTCAACATATGCACGCTGCA |
| 46 | GCCAACATGGATCCTCAGCAGC |
| 47 | GCGCAGATCCGCTAGCTTCT |
| 48 | CCTCGTTTTGGTTGAGGCGG |
| 49 | GCTTTCGTACTGAGATCCGCGATCC |
| 50 | CGCAGGTGGGGGAGGATTCTTGAA |

**Table S2:** Construction of plasmids used in this study. Numbers represent oligonucleotide pairs used for PCR (see Table A1). The restriction enzymes were used for linearization of the vectors and plasmids were assembled using Gibson assembly (multiple primer pairs indicate that constructs were assembled using multiple fragments). To verify the plasmids and the integrations using sequencing, the following primers were used: *accD2:* 21 – 26; *accD3*: 27 -32; *pks*: 33 – 48.

| **Plasmid** | **Template** | **Primers** | **Vector** | **Restriction Enzymes** |
| --- | --- | --- | --- | --- |
| pK19*mobsacB*-*accD2*_G1049A | *C. glutamicum* chromosome | 1 + 2, 3 + 4, 5 + 6, 7 + 8 | pK19*mobsacB* | *BamHI *EcoRI |
| pK19*mobsacB*-*accD3*_G1022A | *C. glutamicum* chromosome | 9 + 10, 11 + 12 | pK19*mobsacB* | *BamHI *EcoRI |
| pK19*mobsacB*-*pks*_T2656G | *C. glutamicum* chromosome | 13 + 14, 15 + 16, 17 + 18, 19 + 20 | pK19*mobsacB* | *BamHI *EcoRI |

**Table S6**. Prediction and (if available) experimental proof of the life style of different corynephages.

| **Phage** | **Domains (temperate)** | **Domains (virulent)** | **Predicted Life Style** | **Experimental** | **Reference** |
| --- | --- | --- | --- | --- | --- |
| CL31 | 5 | 1 | Temperate | - | This work |
| Phi16 | 7 | 3 | Temperate | Temperate | [1,2] |
| Phi673 | 0 | 3 | Virulent | Virulent | [3] |
| Phi674 | 0 | 6 | Virulent | Virulent | [3] |

Figure S1

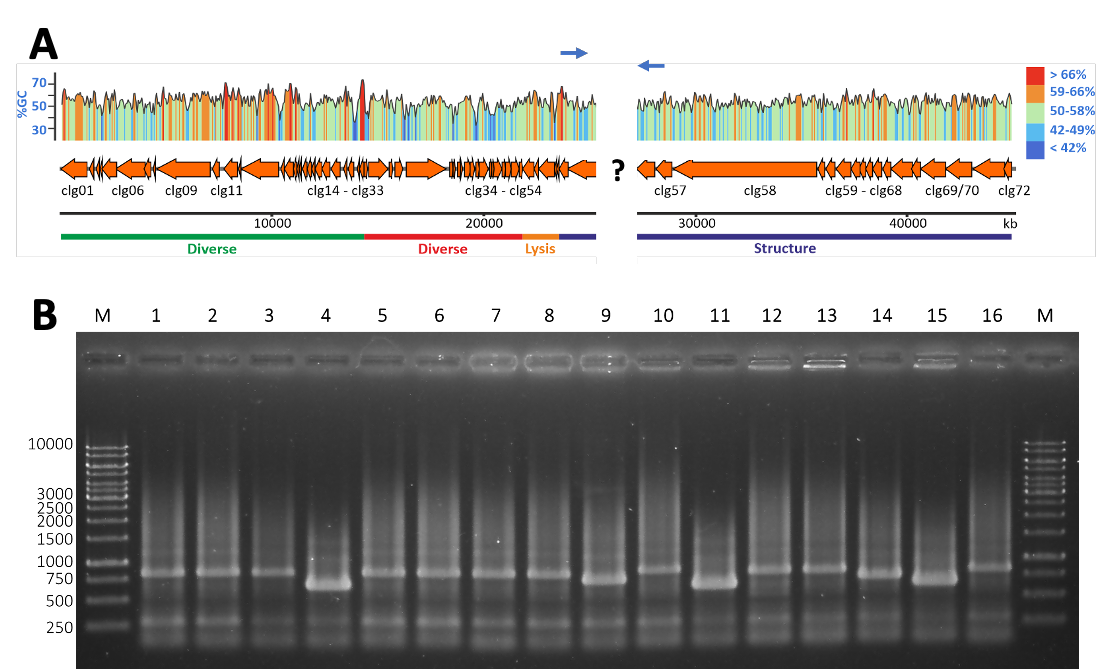

Figure S1: PCR and Sanger sequencing of singles plaques to close the CL31 genome. (A) Schematic representation of the position of a repeat region in the CL31 genome. This region could not be correctly assembled based on Illumina reads and thus was further analyzed with Sanger sequencing. The blue arrows indicate the approximate position of the primers used for PCR amplification and sequencing. (B) Primers 49 and 50 were used to amplify the area shown in A from single CL31 plaques (exemplary plate shown in Figure 2A of the main text). Three different product sizes were obtained. For sequencing three different sizes were chosen (Lane 1, 4, 14).

Figure S2

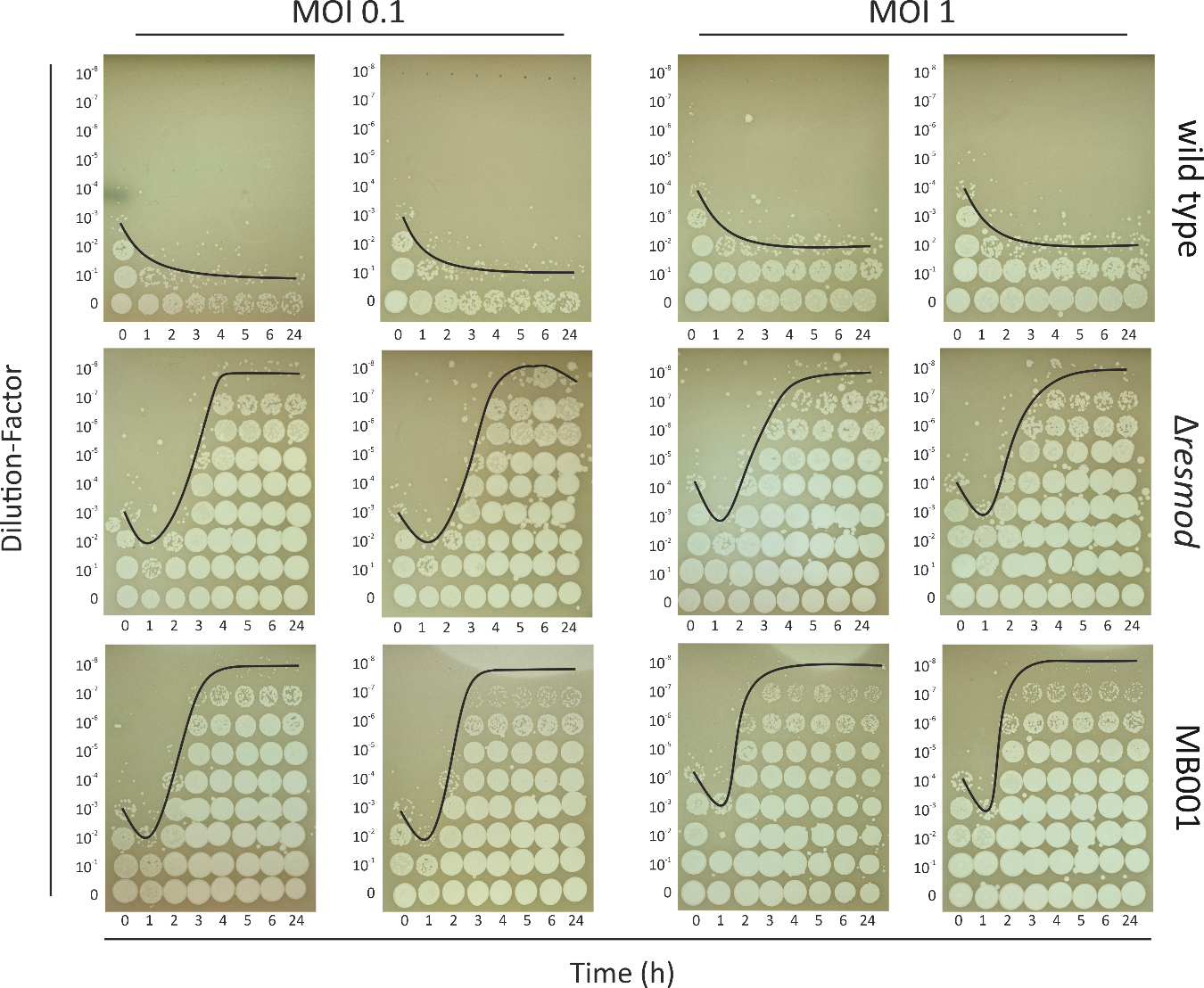

Figure S2: Plate assays as infection dynamics analysis. Infection assay of CL31 with different host strains was used to determine the phage titer at different time points of incubation. For this purpose, bacterial cells were incubated with CL31 in BHI medium with an initial MOI of 0.1 or 1. At the described time points, samples were extracted and 3 µl of the centrifuged supernatant were spotted on a MB001 loan (OD600 0.5).

Figure S3

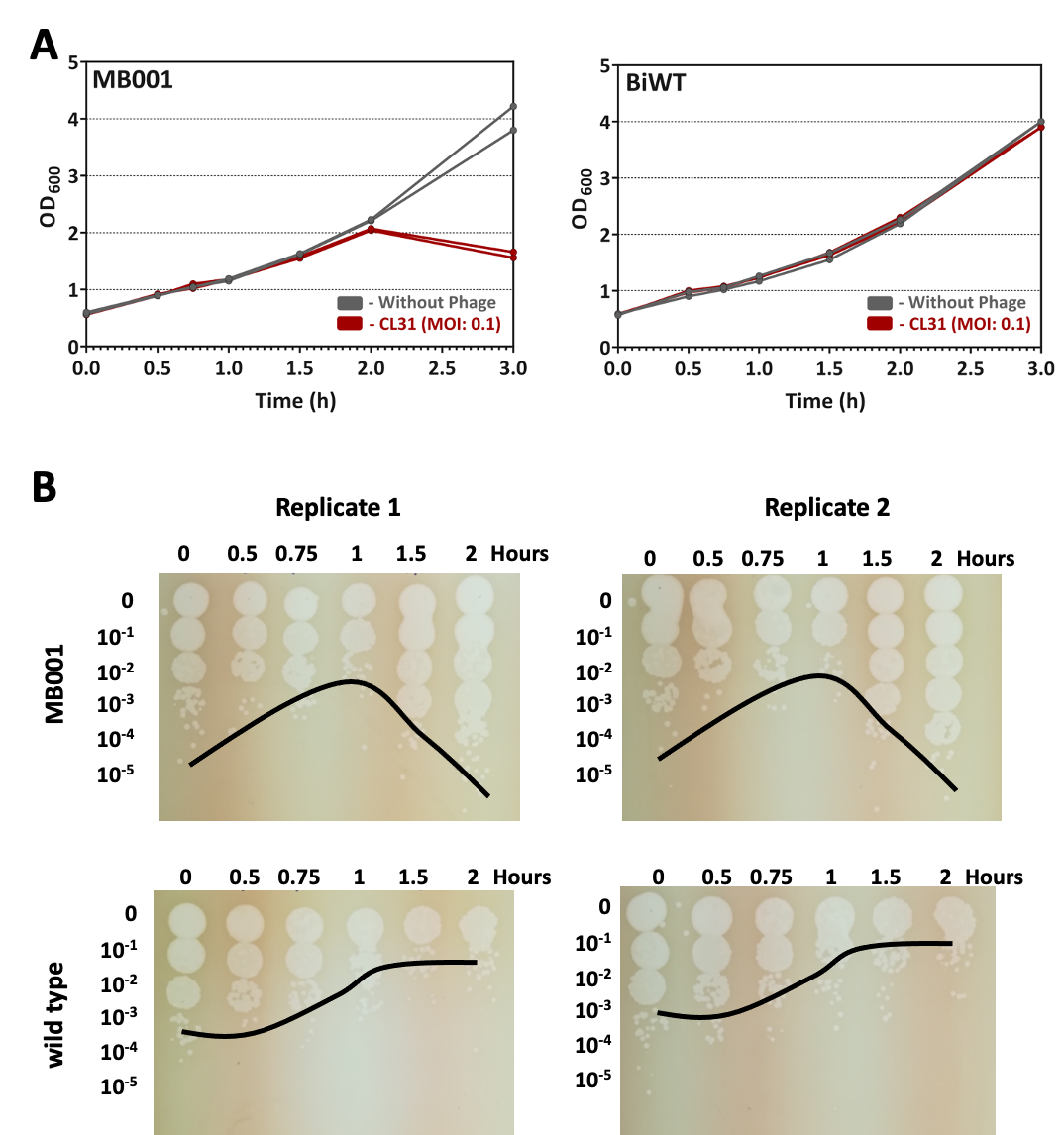

Figure S3: Infection curve for RNA-seq analysis. To determine the best point for RNA-seq analysis, an infection assay of CL31 with wild type or MB001 strains was used to determine the phage titer as well as the growth state (OD_600_ measurements) during incubation. For this purpose, bacterial cells (Starting-OD_600_ 0.5) were incubated with CL31 in BHI medium with an initial MOI of 0.1. At the described time points, samples were extracted, OD measurements were conducted, and 3 µl of the centrifuged supernatant were spotted on a MB001 loan (OD_600_ 0.5). The resulting plates were analysed after an overnight incubation. To cover the moment of highest intracellular phage activity, 1h after cultivation start was chosen to be used for RNA preparation. This represents the point before phage titer increase.
